# MetaBinner: a high-performance and stand-alone ensemble binning method to recover individual genomes from complex microbial communities

**DOI:** 10.1101/2021.07.25.453671

**Authors:** Ziye Wang, Pingqin Huang, Ronghui You, Fengzhu Sun, Shanfeng Zhu

## Abstract

Binning is an essential procedure during metagenomic data analysis. However, the available individual binning methods usually do not simultaneously fully use different features or biological information. Furthermore, it is challenging to integrate multiple binning results efficiently and effectively. Therefore, we developed an ensemble binner, MetaBinner, which generates component results with multiple types of features and utilizes single-copy gene (SCG) information for k-means initialization. It then utilizes a two-step ensemble strategy based on SCGs to integrate the component results. Extensive experimental results over three large-scale simulated datasets and one real-world dataset demonstrate that MetaBinner outperforms other state-of-the-art individual binners and ensemble binners. MetaBinner is freely available at https://github.com/ziyewang/MetaBinner.

## Introduction

Metagenomics, the genomic analysis of microbial communities, provides a culture-independent way for exploring the unknown microbial organisms [1, 2]. Computational methods play an important role in metagenomic studies [3, 4]. Among these computational methods, contig binning aims to put the assembled genomic fragments, contigs, from the same genome into the same bin. The contigs from these bins are then reassembled to form metagenome-assembled genomes (MAG). It is crucial for reconstructing MAGs from metagenomes for further analysis, such as identifying the uncultured bacterial species or viruses [5, 6, 7], associating viruses or bacterium with complex diseases [7, 8, 9] and exploring population diversity [10, 11]. The quality of the MAGs generated by the binners will affect the results of these subsequent analyses. In this paper, we focus on the contig binning methods in general metagenomic data analysis.

Several binning methods have been widely used. CONCOCT [12] is a representative binner that group all the contigs into genomic bins directly. CONCOCT combines coverage vector and tetra-mer frequency vector into one vector for each contig. It uses principal components analysis (PCA) for dimensionality reduction and Gaussian Mixture Model (GMM) for contig binning. MetaBAT 2 [13] is an efficient adaptive binning method that groups some of the contigs whose binning results are the most reliable at first (e.g., the longer contigs) and then gradually add the remaining contigs into the formed genomic bins. MaxBin [14, 15] multiplies the probability *P*_*dist*_ and the probability *P*_*cov*_ that a sequence belongs to a bin based on nucleotide frequency distance and coverage, respectively. A deep learning-based binner, VAMB [16], has recently been developed, which utilizes variational autoencoders (VAE) [17] to convert nucleotide information and coverage information for binning. VAMB then clusters the transformed data using an adaptive iterative medoid method.

Despite the extensive studies, none of the individual binners perform best in all the situations [18]. Therefore, ensemble binning methods are developed to improve the binning performance. The ensemble binning methods can be divided into two categories: 1) the binners that integrate the binning results of other contig binners, such as DAS Tool [19], Binning refiner [20] and MetaWRAP [21], and 2) the stand-alone binners that integrate multiple different component binning results within the ensemble binner, such as BMC3C [22]. DAS Tool [19] realizes genome reconstruction through a dereplication, aggregation, and scoring strategy. It calculates the scores of bins obtained by different binners with bacterial or archaeal reference single-copy genes (rSCG) [23, 24] and chooses the bins with the highest scores. Binning refiner [20] merges results from multiple binning algorithms according to the shared contigs of two bins. The shared contigs between two binners are obtained using BlastN [25]. Then it takes the sets of shared contigs with enough total length as refined bins. MetaWRAP [21] uses Binning refiner [20] to generate hybrid bin sets and chooses the final bins with CheckM [26], which estimates bin quality based on SCGs. UniteM (https://github.com/dparks1134/UniteM) is an ensemble binner developed based on CheckM [26] and Das Tool [19]. Its “greedy” mode uses the SCGs from the bacteria and archaea domain in CheckM to estimate the bin quality and to determine the highest quality MAGs. In contrast, BMC3C [22] is independent of the results from other binners. It repeats k-means clustering multiple times with random initializations to obtain multiple component binning results using the same feature matrix (e.g., 50 times). Then it transforms the index of the results into an affinity matrix. Finally, normalized cut [27] is used for binning. The ensemble methods usually achieve better performance than the individual methods [28].

Although many methods have been proposed to tackle the binning task, a few fundamental issues remain unresolved. Firstly, important biological knowledge such as single-copy genes (SCGs) has largely been ignored in the clustering process by most individual binners and BMC3C. Single-copy marker genes are the genes identified as a single copy in over a certain proportion among the genomes within a specific phylum [29]. Due to this characteristic of the single-copy marker genes, they can be used for estimating the completeness and contamination of the microbial genomes recovered from the metagenomes [26], which enables to evaluate the binning performance without reference genomes and assist the binning process [30, 31]. Secondly, individual binners and BMC3C lack diversity in terms of features. An individual binner usually uses the same features, and BMC3C integrates multiple binning results using the same features. However, various combinations of the features may help reconstruct the complex structure of the metagenomic datasets. The lack of diversities in features also weakens the effectiveness of other ensemble binners that depend on the results from the individual binners. Thirdly, the high-performance ensemble binner, MetaWRAP [21], can only integrate no more than three binning results simultaneously.

Here, we develop a novel ensemble contig binner, MetaBinner, independent of the results from other individual binners. K-means clustering is an efficient clustering method that can be used for large-scale datasets and can produce highly diverse results with different features and initializations [32, 33]. Metabinner first utilizes single-copy gene information for k-means initialization and uses different features for the k-means clustering method to generate different component binning results effectively. It then integrates the component binning results using an efficient two-step ensemble strategy inspired by MetaWRAP [21] and UniteM ‘greedy’ strategy (https://github.com/dparks1134/UniteM). Our binning strategy is designed considering the following aspects. First, we apply k-means clustering as the base clustering method to deal with large-scale datasets. Second, we use single-copy marker gene information for initializations to obtain component binning results with good quality. Next, we use different combinations of the features and different initializations for the k-means clustering to obtain results with diversity for integration. Finally, the two-stage ensemble strategy is applied to select the bins with high completeness and low contamination efficiently and effectively (see “Materials and methods” section for details).

We validated the binning performance of MetaBinner using AMBER [28] and CheckM [26] on three large-scale multi-sample simulated datasets and a real-world dataset. Our experimental results show that MetaBinner outperforms the state-of-the-art binners, including CONCOCT, MetaBAT, MaxBin, VAMB, DAS Tool, MetaWRAP and BMC3C. For the simulated datasets, MetaBinner increases 46.7% and 20.4% on average in terms of the numbers of the near-complete bins (> 90% completeness and < 5 % contamination), compared with the best individual binner and the second-best ensemble binner, respectively.

## Results

### MetaBinner: a novel ensemble method for contig binning

MetaBinner has five major steps. (i) Construct the feature representations of contigs with coverage and composition information; (ii) Determine the number of bins; (iii) Generate binning results with multiple features and initializations; (iv) Split bins with high contamination and completeness according to the single-copy marker genes; (v) Incorporate the component binning results with a two-step efficient ensemble strategy. The general pipeline of MetaBinner is shown in Figure 1. Detailed explanations of the steps are given in the “Materials and methods” section.

**Figure 1.**
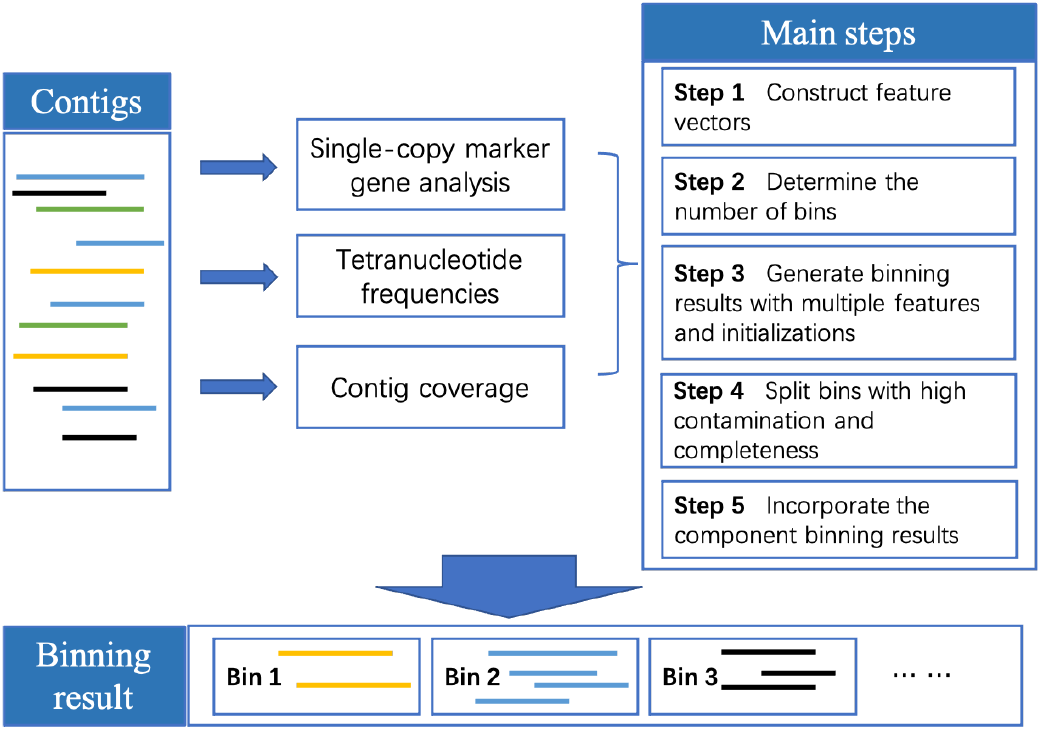
The general workflow of MetaBinner for contig binning.

In the following, we compared the performance of Metabinner with other individual binners (CONCOCT [12], MetaBAT [34, 13], MaxBin [14, 15], VAMB [16]) and ensemble binners (BMC3C [22], MetaWRAP [21] and DAS Tool [19]). Then, we conducted experiments to show the necessity and effectiveness of multiple features and initializations. Finally, we showed the running time of the binners on several datasets.

### MetaBinner outperforms other available contig binning methods on the simulated datasets evaluated by AMBER [28]

We used three large-scale simulated multi-sample datasets to evaluate the binners using the evaluation metrics proposed in AMBER [28]. Detailed explanations of the datasets and evaluation metrics are given in the “Materials and methods” section. Table 1 shows that MetaBinner can recover much more high-quality genomes than other binners under different completeness and contamination thresholds. None of the four individual binners perform best in all the three simulated datasets in terms of the number of high-quality bins, supporting the similar statement given in [18]. MetaBAT achieves good performance on these simulated datasets among the individual binners. BMC3C estimates the bin number based on the contig numbers, which may affect its performance. The total number of predicted bins per binner and the true bin number for each dataset are available as Appendix Table A1.

**Table 1.** Performance comparison of the binners on the simulated datasets evaluated by AMBER.

| Dataset | Methods | Metrics |  |  |  |  |  |
| --- | --- | --- | --- | --- | --- | --- | --- |
|  |  | #bins<br>(>50%<br>comp<br><10%<br>cont) | #bins<br>(>70%<br>comp<br><10%<br>cont) | #bins<br>(>90%<br>comp<br><10%<br>cont) | #bins<br>(>50%<br>comp<br><5%<br>cont) | #bins<br>(>70%<br>comp<br><5%<br>cont) | #bins<br>(>90%<br>comp<br><5%<br>cont) |
| CAMI Airways | CONCOCT | 42 | 38 | 32 | 42 | 38 | 32 |
|  | MaxBin | 88 | 82 | 64 | 67 | 62 | 51 |
|  | MetaBAT | 92 | 80 | 56 | 84 | 74 | 51 |
|  | VAMB | 90 | 75 | 47 | 86 | 72 | 46 |
|  | BMC3C | 18 | 8 | 6 | 13 | 6 | 4 |
|  | DAS Tool | 106 | 101 | 80 | 94 | 91 | 71 |
|  | MetaWRAP | 136 | 119 | 83 | 131 | 118 | 82 |
|  | MetaBinner | **215** | **191** | **144** | **186** | **169** | **129** |
| CAMI<br>Gastrointestinal<br>tract | CONCOCT | 66 | 64 | 61 | 62 | 60 | 57 |
|  | MaxBin | 114 | 110 | 101 | 106 | 103 | 97 |
|  | MetaBAT | 97 | 93 | 83 | 93 | 89 | 80 |
|  | VAMB | 57 | 55 | 42 | 56 | 55 | 42 |
|  | BMC3C | 12 | 9 | 7 | 7 | 4 | 3 |
|  | DAS Tool | 130 | 125 | 116 | 124 | 120 | 111 |
|  | MetaWRAP | 134 | 126 | 114 | 131 | 123 | 112 |
|  | MetaBinner | **183** | **173** | **152** | **176** | **166** | **147** |
| CAMI mouse gut | CONCOCT* | 95 | 95 | 84 | 88 | 88 | 79 |
|  | MaxBin* | 439 | 419 | 342 | 401 | 386 | 319 |
|  | MetaBAT* | 353 | 318 | 240 | 339 | 309 | 236 |
|  | VAMB | 382 | 370 | 293 | 372 | 363 | 288 |
|  | BMC3C | 315 | 306 | 262 | 297 | 290 | 253 |
|  | DAS Tool* | 460 | 449 | 354 | 422 | 416 | 334 |
|  | MetaWRAP | 506 | 487 | 377 | 500 | 482 | 375 |
|  | MetaBinner | **575** | **532** | **421** | **535** | **503** | **409** |

The best results among all the methods are in bold, while the best results among the individual binners are italicized. The input binning results of MetaWRAP and DAS Tool are generated by CONCOCT, MaxBin and MetaBAT. “#bins (>50% comp <10% cont)” denotes that the number of recovered bins that have >50% completeness and <10% contamination. The results with “\*” are from the CAMI tutorial [4].

**Table 2.**
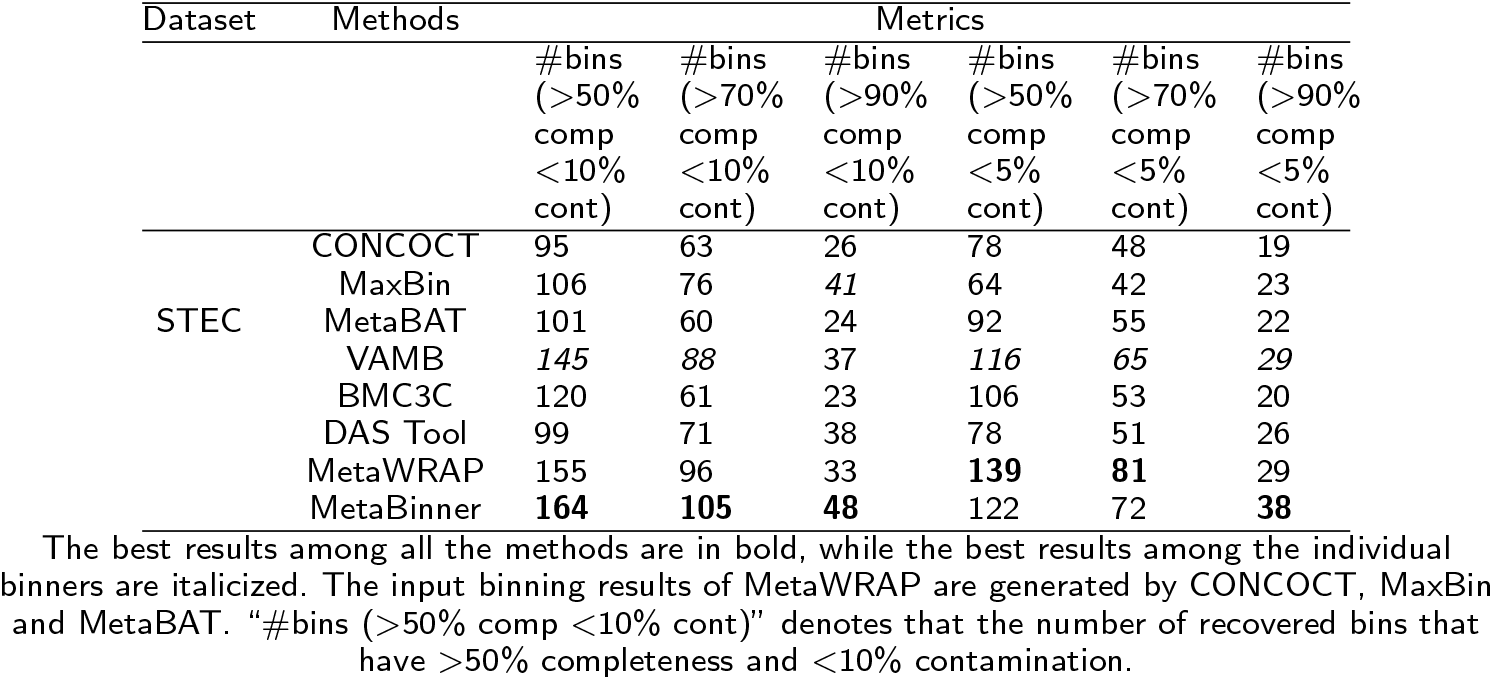
Performance comparison of the binners on the real dataset evaluated by CheckM.

Take the CAMI Gastrointestinal tract as an example for analysis. From the experimental results, we have three main findings. First, MetaBinner recovered the most high-quality genomes (>50% completeness and <10% contamination). Specifically, compared with the second-best binner, MetaBinner improves the numbers of near-complete (NC) genomes (> 90% completeness and < 5 % contamination; as defined in VAMB) from 112 to 147. Secondly, BMC3C, MetaBAT and CONCOCT assign the most base pairs at the cost of a lower Adjusted Rand Index (ARI) (Fig. 2a). Among the binners with the highest ARI (over 90%), MetaBinner assigns the most base pairs. Thirdly, among all the binners, MetaBinner achieves the highest average completeness of all predicted bins. Its average purity is close to those of the binners with the highest average purity (Fig. 2b).

**Figure 2.**
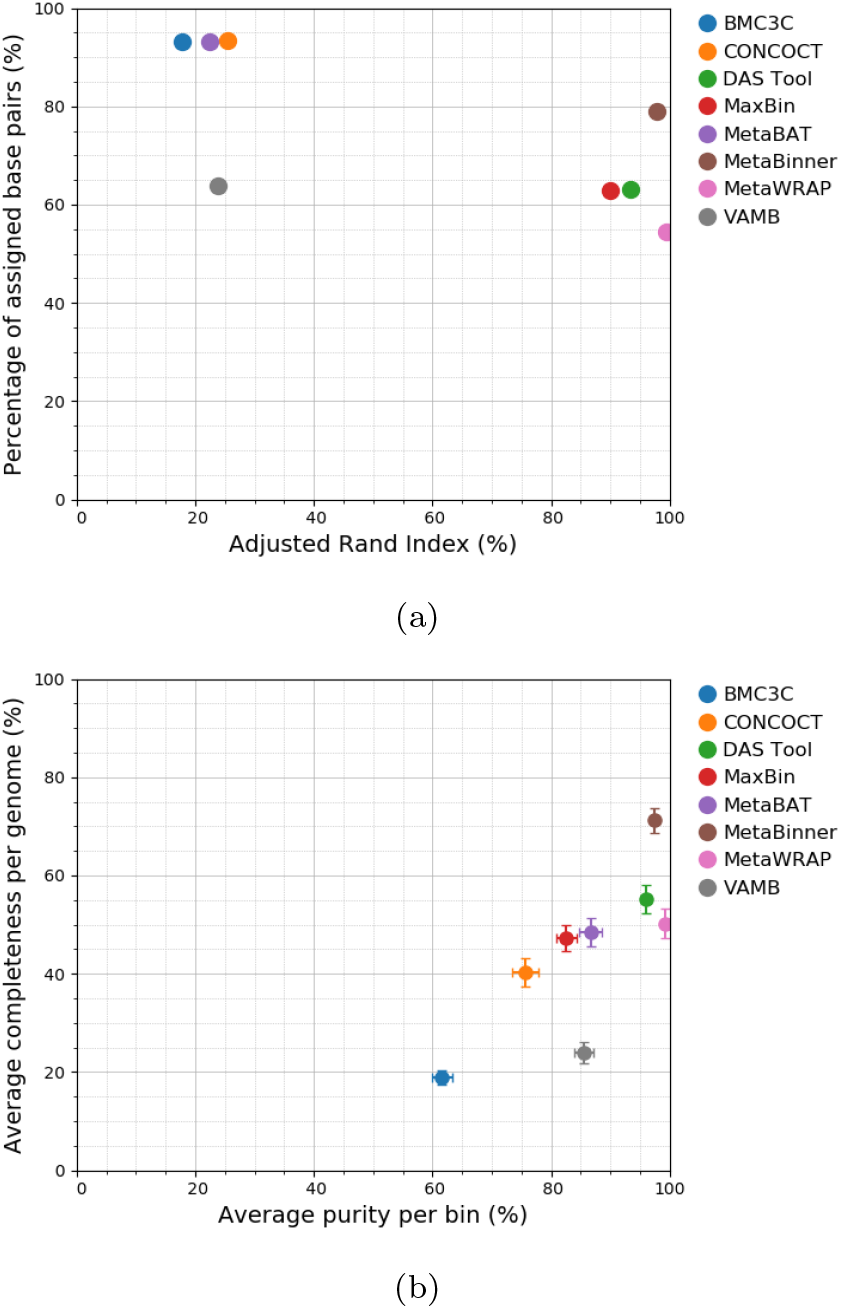
Assessing binners on the CAMI Gastrointestinal tract dataset based on base pairs. a) Adjusted Rand index (x axis) versus percentage of assigned base pairs (y axis). b) Average purity (x axis) versus average completeness (y axis) of all predicted bins per binner.

### MetaBinner produces the most high-quality MAGs on the real dataset

For the real dataset, the true genomes in the metagenomes are unknown. In this situation, CheckM is widely used in studies [35, 36] for selecting the high quality bins from the Metagenome-Assembled Genomes (MAGs). As shown in Table A2, MetaBinner and MetaWRAP achieve the best overall performance. Among the individual binners, VAMB achieves the best performance. As shown in the table, MetaBinner can recover the most near-complete genomes. The assembly results of the sequencing data may affect the follow-up binning performance. Therefore, we assembled the reads using another popular assembler, MetaSPAdes, and used the assembled contigs for binning. The results are given in Appendix Table A2, and MetaBinner also achieves the best performance.

### The effect of generating component binning results with multiple features and initializations

When running the k-means++-based method for comparison, we use the length of each contig to set the weight for each contig. We take the binning results of CAMI Airways as the example.

#### 1) The effect of the “Partial Seed” method

To demonstrate the effect of the “Partial seed” method, we ran k-means++ randomly for three times and compared the results with three “Partial seed” binning results for each feature matrix generated by “Step 3”. We suppose that the improvement of binning results is not only because we used contigs containing single-copy marker genes as cluster centers but also because we added the regular k-means++ initialization part. To further prove this point, we also compared the “partial seed” results with the results using the same cluster centers but without the regular k-means++ initialization part. “Seed k-means” indicates running k-means++ using the features of the contigs containing the chosen single-copy marker gene as the cluster centers, and the bin number is the same as the number of corresponding contigs.

From the experimental results given in Table 3, we have the following main findings. Firstly, the component binning results generated using *X*_*combo*_ feature matrix (109 high-quality bins) have the best performance compared with those using other feature matrices (41 and 93 high-quality bins). Secondly, the length-weight strategy can improve the binning performance. Thirdly, using the contigs containing the single-copy marker genes as the cluster centers can improve the binning performance. Take the results using *X*_*combo*_ feature matrix as an example. The “seed k-means” method recovers about 100 high-quality bins on average, compared with 75.33 generated by regular k-means++. Finally, the part with regular k-means++ initialization in “Partial seed” helps in binning, especially for the component binning results generated using *X*_*combo*_ and *X*_*cov*_ feature matrix.

**Table 3.** Performance comparison of “Partial seed” and regular k-means++ in terms of recovered high-quality bins in CAMI Airways dataset.

| Feature | Methods | Metrics |  |  |  |  |  |
| --- | --- | --- | --- | --- | --- | --- | --- |
|  |  | #bins<br>(>50%<br>comp<br><10%<br>cont) | #bins<br>(>70%<br>comp<br><10%<br>cont) | #bins<br>(>90%<br>comp<br><10%<br>cont) | #bins<br>(>50%<br>comp<br><5%<br>cont) | #bins<br>(>70%<br>comp<br><5%<br>cont) | #bins<br>(>90%<br>comp<br><5%<br>cont) |
| $X_{combo}$ | k-means++ average (no length weighting) | 32.67 | 26.33 | 9.33 | 27.67 | 21.67 | 6.67 |
|  | k-means++ average | 75.33 | 60.33 | 39.00 | 65.67 | 52.67 | 35.33 |
|  | seed k-means average | 100.33 | 91.33 | 63.00 | 74.67 | 67.33 | 49.00 |
|  | Partial seed average | **109.33** | **96.00** | **65.67** | **81.67** | **71.33** | **50.67** |
| $X_{cov}$ | k-means++ average (no length weighting) | 30.67 | 26.67 | 20.33 | 18.00 | 16.00 | 11.33 |
|  | k-means++ average | 57.00 | 51.00 | 45.33 | 43.67 | 38.67 | 35.00 |
|  | seed k-means average | 89.00 | 82.67 | 75.00 | 66.00 | 61.33 | 57.00 |
|  | Partial seed average | **93.00** | **86.33** | **78.67** | **71.00** | **66.33** | **61.00** |
| $X_{com}$ | k-means++ average (no length weighting) | 14.33 | 7.33 | 1.00 | 11.33 | 7.00 | 1.00 |
|  | K-mean++ average | 30.33 | 22.67 | 13.33 | 21.00 | 16.00 | 7.33 |
|  | seed k-means average | 40.67 | **29.67** | 15.67 | **24.00** | 16.00 | 7.67 |
|  | Partial seed average | **41.00** | 29.00 | **16.67** | 23.67 | **16.33** | **8.67** |

The best results based on each feature matrix are in bold. $X_{combo}$ denotes the feature matrix combining coverage and composition information, $X_{cov}$ denotes the feature matrix using coverage information, and $X_{com}$ denotes the feature matrix using composition information (see “Materials and methods” for more details). “#bins (>50% comp <10% cont)” denotes that the number of recovered bins that have >50% completeness and <10% contamination. “no length weighting” denotes that all contigs are assigned equal weight while running k-means++.

#### 2) The effect of incorporating binning results using three kinds of feature combinations

We use the changes of the final output of MetaBinner to reflect the effect. Metabin_A, Metabin_B and Metabin_C denote the ensemble results of the component binning results generated using *X*_*combo*_, *X*_*cov*_, and *X*_*com*_, respectively. Metabin_AB deontes the MetaBinner results after removing the parts related to Metabin_C (see Figure 3) during the second step of integration. The results drop from 215 to 203, 197, and 189 after removing the components using each feature matrix in terms of the number of high-quality bins (Table 4). Interestingly, the Metabin_A (175 high-quality bins) even has better performance than the second-best method (MetaWRAP: 136 high-quality bins) for this dataset (Table 4).

**Figure 3.**
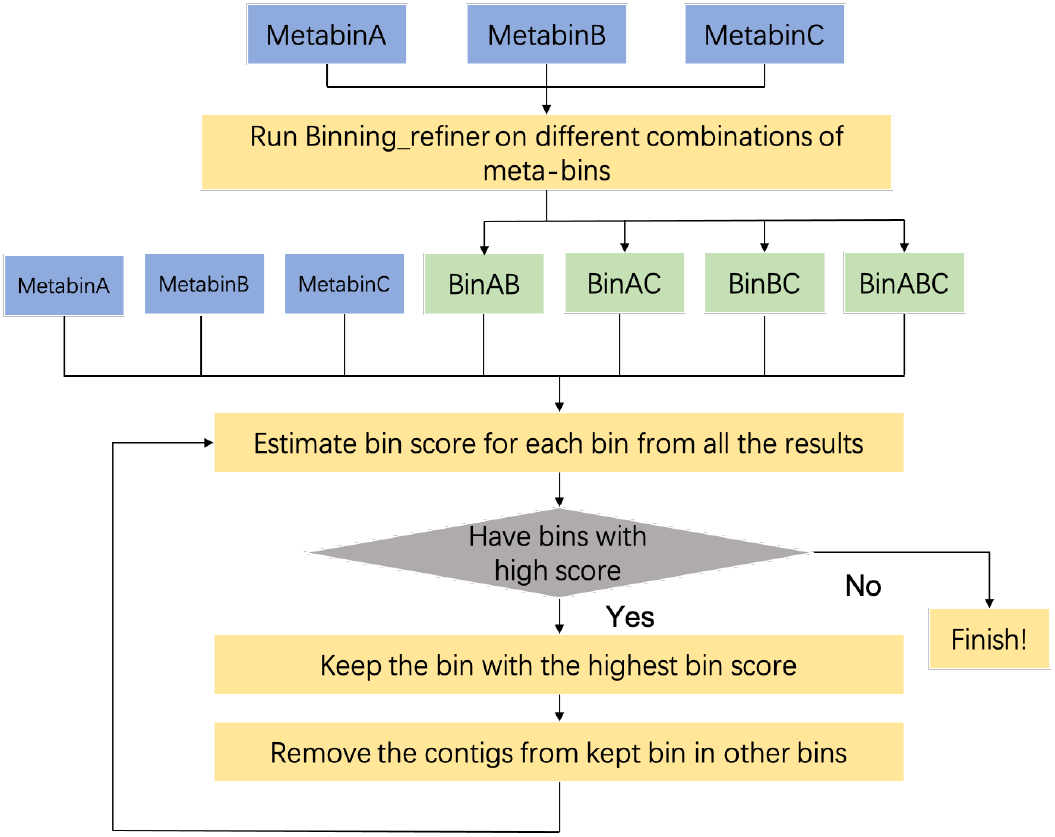
The ensemble strategy workflow (the second stage). BinAB denotes the refined binning results of MetabinA and MetabinB using Binning refiner [20].

**Table 4.** Performance comparison of MetaBinner and the results of MetaBinner after removing binning results using one or two kinds of feature combinations in CAMI Airways dataset.

| Methods | Metrics |  |  |  |  |  |
| --- | --- | --- | --- | --- | --- | --- |
|  | #bins<br>(>50%<br>comp<br><10%<br>cont) | #bins<br>(>70%<br>comp<br><10%<br>cont) | #bins<br>(>90%<br>comp<br><10%<br>cont) | #bins<br>(>50%<br>comp<br><5%<br>cont) | #bins<br>(>70%<br>comp<br><5%<br>cont) | #bins<br>(>90%<br>comp<br><5%<br>cont) |
| Metabin_A | 175 | 157 | 113 | 130 | 120 | 87 |
| Metabin_B | 145 | 139 | 116 | 113 | 110 | 89 |
| Metabin_C | 148 | 119 | 96 | 114 | 92 | 73 |
| Metabin_AB | 203 | 184 | 134 | 163 | 151 | 111 |
| Metabin_AC | 197 | 173 | 130 | 167 | 151 | 116 |
| Metabin_BC | 189 | 166 | 135 | 163 | 146 | 118 |
| Metabin_ABC (MetaBinner) | **215** | **191** | **144** | **186** | **169** | **129** |

The best results are in bold. “#bins (>50% comp <10% cont)” denotes that the number of recovered bins that have >50% completeness and <10% contamination.

#### 3) The effect of integrating component results using the proposed ensemble strat egy instead of DAS Tool

Metabin A (DAS Tool), Metabin B (DAS Tool) and Metabin C (DAS Tool) denote the DAS Tool integration results of the component binning results generated using *X*_*combo*_, *X*_*cov*_, and *X*_*com*_, respectively. Table 5 shows the performance comparison of MetaBinner and DAS Tool using one feature combination in CAMI Airways dataset. The proposed ensemble strategy has better performance than DAS Tool on all the three feature combinations. The number of high quality bins using *X*_*combo*_ feature matrix improves from 147 to 175.

**Table 5.** Performance comparison the results of MetaBinner and DAS Tool using one feature combination in CAMI Airways dataset.

| Methods | Metrics |  |  |  |  |  |
| --- | --- | --- | --- | --- | --- | --- |
|  | #bins<br>(>50%<br>comp<br><10%<br>cont) | #bins<br>(>70%<br>comp<br><10%<br>cont) | #bins<br>(>90%<br>comp<br><10%<br>cont) | #bins<br>(>50%<br>comp<br><5%<br>cont) | #bins<br>(>70%<br>comp<br><5%<br>cont) | #bins<br>(>90%<br>comp<br><5%<br>cont) |
| Metabin_A | **175** | **157** | **113** | **130** | **120** | **87** |
| Metabin_A (DAS Tool) | 147 | 138 | 106 | 108 | 103 | 80 |
| Metabin_B | **145** | **139** | **116** | **113** | **110** | **89** |
| Metabin_B (DAS Tool) | 131 | 127 | 112 | 100 | 98 | 84 |
| Metabin_C | **148** | **119** | **96** | **114** | **92** | **73** |
| Metabin_C (DAS Tool) | 112 | 106 | 93 | 86 | 81 | 71 |

The best results based on each feature matrix are in bold. “#bins (>50% comp <10% cont)” denotes that the number of recovered bins that have >50% completeness and <10% contamination.

### Running time of the binners

All the results given in this section were run on two Intel Xeon CPUs (E5-2660 v3, 2.60GHz) with 128G RAM. We ran all the binners with multiple threads. Table 6 shows the running time of MetaBinner, the three individual binners for DAS Tool and MetaWRAP and the two ensemble binners on different datasets (CAMI Airways and CAMI mouse gut). Since most binners can share the steps of generating composition and coverage files, we only compared the running time required after generating these files. For the dataset with more samples (CAMI mouse gut), it takes much more time to run MaxBin (more than 7,100 minutes) compared with other binners, so the whole running time of MetaBinner (about 1,514 minutes) is much less than other ensemble binners. For the CAMI airways dataset, the running time of MetaBinner is similar to that of DAS Tool.

**Table 6.** The running time the binners.

| Binners | CAMI Airways | CAMI mouse gut |
| --- | --- | --- |
| CONCOCT | 39m21s | 166m40s |
| MaxBin | 933m51s | 7120m52s |
| MetaBAT | **22m58s** | **16m12s** |
| DAS Tool | 19m47s | 25m36s |
| MetaWRAP | 547m15s | 1601m26s |
| DAS Tool (+running time of three individual binners) | **1015m57s** | 7320m20s |
| MetaWRAP (+running time of three individual binners) | 1543m25s | 8905m10s |
| MetaBinner | 1057m48s | **1514m24s** |

The best results among the individual binners and the ensemble binners are in bold.

## Discussion

In this paper, we introduced MetaBinner, a novel stand-alone ensemble binner for large-scale contig binning. Firstly, MetaBinner generates binning results mainly using “Partial seed” k-means with multiple types of features and initializations. Then, MetaBinner applies an effective and efficient two-step ensemble strategy to integrate the component binning results. We compared MetaBinner with advanced binning tools, CONCOCT, MaxBin, MetaBAT, VAMB, BMC3C, DAS Tool, and MetaWRAP on four datasets, and MetaBinner has the best overall performance among all the datasets. We also show the effect of the “Partial seed” method proposed in the paper and the impact of multiple features and initializations.

MetaBinner applies single-copy marker genes in the clustering process by the “Partial seed” k-means method and solves the problem that the approximate number estimation may affect the performance of the methods based on single-copy marker genes. The approximate number estimation may be caused by the contigs from other categories of the taxon in complex metagenomic communities or the imperfect metagenomics assembly [4]. Secondly, MetaBinner improves the binning performance by generating component results using different feature matrices and integrating the component results with a two-step ensemble strategy based on SCGs. The above points make up for the shortcomings of the existing binning methods. Furthermore, MetaBinner is a stand-alone ensemble binner, which doesn’t utilize the results from other individual binners as the other two popular ensemble binners, MetaWRAP and DAS Tool, which may reduce running time. MetaBinner can be integrated into different ensemble approaches (such as MetaWRAP and DAS Tool) as a component to achieve better performance. Other individual binners can also be integrated into MetaBinner flexibly by replacing Metabin A, B, or C with their results.

Despite the successes of the MetaBinner for large-scale contig binning, it still has limitations. For example, the single-copy marker gene sets only contain the bacterial and archaeal reference genes. In the future, we would like to explore the way to integrate the marker genes from other taxa, such as microbial eukaryotes [37], into the binning pipeline to resolve more complex microbial communities. Furthermore, since some component binning results for integration are generated using coverage information alone as features, we recommend applying MetaBinner to multi-sample datasets.

## Conclusion

MetaBinner provides a powerful method for large-scale contig binning. We evaluated MetaBinner and the compared methods based on real and simulated datasets. The results show that MetaBinner outperforms other individual binners, BMC3C, and the ensemble binners based on the results from multiple individual binners.

## Materials and methods

In this section, we present: 1) the descriptions of the benchmark datasets; 2) the details of each step in MetaBinner; 3) the metrics to evaluate the binning performance; and 4) implementation and parameter settings of different methods.

### Datasets

#### The simulated datasets

We used one benchmark dataset, CAMI mouse gut, from the recent CAMI benchmarking toolkit tutorial [4] and two other ‘toy’ human short-read datasets from CAMI II (https://data.cami-challenge.org), CAMI Airways and CAMI Gastrointestinal tract, to evaluate the performance of the binners. Most competing methods can only cluster the contigs longer than 1,000 bp. Therefore, we kept the contigs of the gold standard cross-sample assembly longer than 1,000 bp for binning. We used the simulated Illumina HiSeq reads that CAMI provided to generate the coverage information. Table 7 shows the general information of the simulated datasets.

**Table 7.** Datasets used in the experiments.

| Data Type | Dataset | Number of samples | Number of contigs (> 1000bp) |
| --- | --- | --- | --- |
| Simulated | CAMI Airways | 10 | 285047 |
|  | CAMI Gastrointestinal tract | 10 | 57088 |
|  | CAMI Mouse gut | 64 | 241451 |
| Real | STEC | 53 | 255484 |

#### The real datasets

To assess the performance of the binners on large real datasets, we used a real dataset with multiple samples, the “STEC” dataset. The “STEC” dataset [38] contains 53 samples from a set of fecal specimens in the PRJEB1775 study (https://www.ebi.ac.uk). MetaWRAP [21] is a modular pipeline, and its “Assembly” module allows users to assemble metagenomic reads with metaSPAdes [39] or MEGAHIT [40]. The reads from all the samples are co-assembled by MetaWRAP-Assembly module with default parameters and the default assembler (MEGAHIT). The general information of the real dataset is given in Table 7.

### The MetaBinner algorithm

Figure 1 shows the framework of MetaBinner, which consists of two modules: 1) “Component module” includes steps 1-4, developed for generating high-quality, diverse component binning results; and 2) “Ensemble module” includes step 5, developed for recovering individual genomes from the component binning results. More descriptions of each step are as follows.

#### Step 1: Construct feature vectors for metagenomic contigs

Each contig co-assembled from *M* samples can be represented with the combination of a coverage vector (*M* dimensional) and a composition vector (*T* dimensional) as done in previous studies [41, 42], where *T* is the number of distinct tetramers. The coverage vector and the composition vector denote the coverage profiles across the *M* samples and the tetramer frequency, respectively. Similar to COCACOLA [41], a small value is added to each entry of the vectors (0.01 for the coverage vector; one for the composition vector) to handle zero values. Then the coverage matrix and the composition matrix are normalized as in [41]. For some datasets with high-quality sequencing with a large number of sequencing samples, the coverage vector contains much more information for binning. In such cases, each contig can be represented by the *M* dimensional coverage vector only. Furthermore, different organisms usually have different tetra-mer composition profiles [43, 44]. Therefore, the feature matrix of the contigs is denoted as *X*_*combo*_ ∈ ℝ^*N×*(*M* +*T*)^, *X*_*cov*_∈ ℝ^*N×M*^ or *X*_*com*_∈ ℝ^*N×T*^, where *N* denotes the number of contigs. We did log transformation for each feature matrix as done in CONCOCT for the *X*_*combo*_ feature matrix. In this way, we obtained three feature matrices for each dataset.

#### Step 2: Determine the number of bins

Similar to SolidBin [42] and COCACOLA [41], we utilized the set of single-copy marker genes universal for bacteria and archaea provided by [14] to estimate the number of genomes in the metagenomic data. As stated in [14], some marker genes may be fragmented into pieces, influencing the estimation of the bin number. So we calculated the number of contigs containing each marker gene, and then used the third quartile value of the numbers in ascending order to determine the initial bin number *k*_0_. A list of numbers larger than *k*_0_ were then sequentially tried as the bin numbers in the k-means algorithm (see Figure S1). The bin number yielding the largest silhouette coefficient [45] value of the binning result is chosen as the final bin number.

#### Step 3: Generate binning results with multiple features and initializations

We proposed the “Partial Seed” strategy based on k-means++ [46], a variant of k-means, to generate high-quality diverse component binning results. More descriptions about k-means++ are given in Appendix.

MaxBin [14, 15] initializes the features of each putative genome (bin) for an expectation-maximization algorithm using tetranucleotide frequencies and coverage levels of the contigs harboring a certain single-copy marker gene. However, the single-copy marker gene sets do not cover all the genomes in the microbial communities. In their method, the bin number equals the number of the contigs containing the certain SCG. In this way, some contigs from the genomes without the certain SCG may be assigned into the wrong bins, resulting in high contamination. Therefore, we proposed the “Partial Seed” strategy. Instead of choosing the centers randomly, we chose a single-copy marker gene and used it to define cluster centers. For a particular single-copy marker gene, suppose that there are *l* contigs containing this gene. These *l* contigs should belong to different genomes. Therefore we used features of these *l* contigs as the initial centers, and generate the other *K* − *l* initial clustering centers by k-means++. *K* denotes the bin number estimated by “Step 2” and is always larger than *k*_0_ mentioned in “Step 2”.

To achieve a better integration effect while using the ensemble module, we can produce several diverse binning results using different sets of fixed initial clustering centers. Therefore, we kept the first, second, and third quartile values of the numbers of contigs containing each marker gene. Similar to MaxBin, the shortest marker gene that corresponds to each number was selected. In this way, we obtained three sets of designated initial clustering centers for each feature matrix.

To integrate the binning results according to the data features themselves without considering fixed initial clustering centers into the final result, we also run regular k-means++ using each feature matrix to generate the basic clustering result. In summary, four binning results are generated for each feature matrix; three of them are from the “Partial Seed” method. Furthermore, there are three feature matrices (*X*_*combo*_, *X*_*cov*_, and *X*_*com*_) for each dataset.

#### Step 4: Split bins with high contamination and completeness according to the single-copy marker genes

For each binning result generated in “Step 3”, we used the SCG sets for the bacteria and archaea domain used by CheckM to estimate the bins’ contamination and completeness. We ran CheckM for one binning result and obtained the contigs having the single-copy marker genes. Similar to UniteM, we then used the information to estimate each bin’s contamination and completeness of each component binning result using the scoring strategy in CheckM. BinSanity [31] applies a composition-based refinement to handle the highly contaminated or low completion bins. In our paper, if a bin has high contamination (>= 50%) and completeness (>= 70%), we split it by estimating the number of sub-bins of the bin using the same approach as in “Step 2” and running the “Partial Seed” strategy using a certain single-copy marker gene. We regard the binning results generated by “Step 4” as “component binning results”.

#### Step 5: Incorporate the component binning results with an ensemble module

The quality of binning results obtained by different input matrices may be markedly different. Therefore, we completed the integration process in two stages. In the first stage of the ensemble module, we separately integrated four component binning results generated by “Step 4” for each input feature matrix. First, the bins with high bin scores estimated by the SCGs for bacterial and archaeal genomes will be selected. Then, the contigs in the selected bins will be removed from other bins. The above two operations are repeated until there are no high-quality bins. In the second stage, we used a method similar to MetaWRAP to integrate the three ensemble results of the first stage, as shown in Figure 3. First, apply Binning refiner to refine bins. Then, use the same method as the first stage to select high-quality bins.

The process of picking high-quality bins for the two stages is highly similar to UniteM’s greedy strategy. The main difference is that MetaBinner bins all the contigs while generating the component binning results. Therefore, we only ran CheckM once for each domain (bacteria and archaea domains) to get the SCG information for all the contigs, instead of running it for all the component results as done in UniteM. For the same reason, the second stage does not need to run CheckM as many times as MetaWRAP.

### Evaluation metrics

For the simulated datasets, we use AMBER [28], which implements the metrics in the first CAMI binning challenge for evaluation [3, 4]. AMBER metrics are calculated based on a gold standard mapping result of the contigs or reads, so it is only suitable for the simulated datasets. We will use the following quantities to evaluate the binning results: a) Number of high-quality genomes; b) Adjusted Rand index (x axis) versus percentage of assigned base pairs (y axis), and c) Average purity (x axis) versus average completeness (y axis) of all predicted bins per method. The definitions of the criteria are given in AMBER [28]. The high-quality genomes are defined as genomes with > 50% completeness and < 10% contamination as done in [4].

For the real dataset without known genome assignments, we applied CheckM [26] for evaluation to obtain the bin’s completeness and contamination scores.

### Implementation and parameter settings

We compared Metabinner with seven advanced binners: CONCOCT-1.0.0, MaxBin 2.2.6, MetaBAT 2.12.1, VAMB 3.0.2, MetaWRAP 1.2.1, DAS Tool 1.1.2 and BMC3C, respectively. MetaWRAP and DAS Tool need to integrate the results from other binners and CONCOCT, MaxBin, and MetaBAT were chosen for these two ensemble binners as done in MetaWRAP [21]. We ran CONCOCT-1.0.0, MaxBin 2.2.6, and MetaBAT 2.12.1 using the binning module of MetaWRAP with the “– universal” parameter to use universal marker genes, which can improve binning for the Archaea genomes. We ran MetaWRAP with “-c 50” to set the minimum % completion of the bins. For a fair comparison with other methods, we ran coassembly mode of VAMB with “–jgi /path/to/depth/depth.txt –minfasta 200000”. The coverage profiles of the contigs were obtained via MetaWRAP 1.2.1 script: “binning.sh”. The results of the simulated and real datasets were evaluated by AMBER 2.0.21-beta and CheckM v1.1.3.

MetaWRAP is a modular pipeline, and we regard its “bin refinement” module as MetaWRAP in this paper if there is no additional explanation.

## Appendix Brief description of of k-means++

K-means++ [46] is a variant of k-means, which utilizes a smart way to initialize clustering centers to improve clustering accuracy and computational speed. K-means++ uniformly chooses a data point as the first initial center *c*_1_ at random from the set of data points, *χ*. A new center *c*_*i*_ is chosen from *χ* with probability *P* (*x*), which is defined in equation (1). The process is repeated until *K* centers have been chosen.

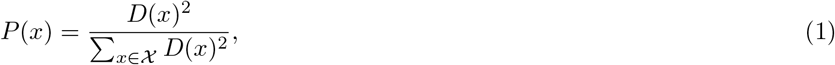

**Figure A1.**
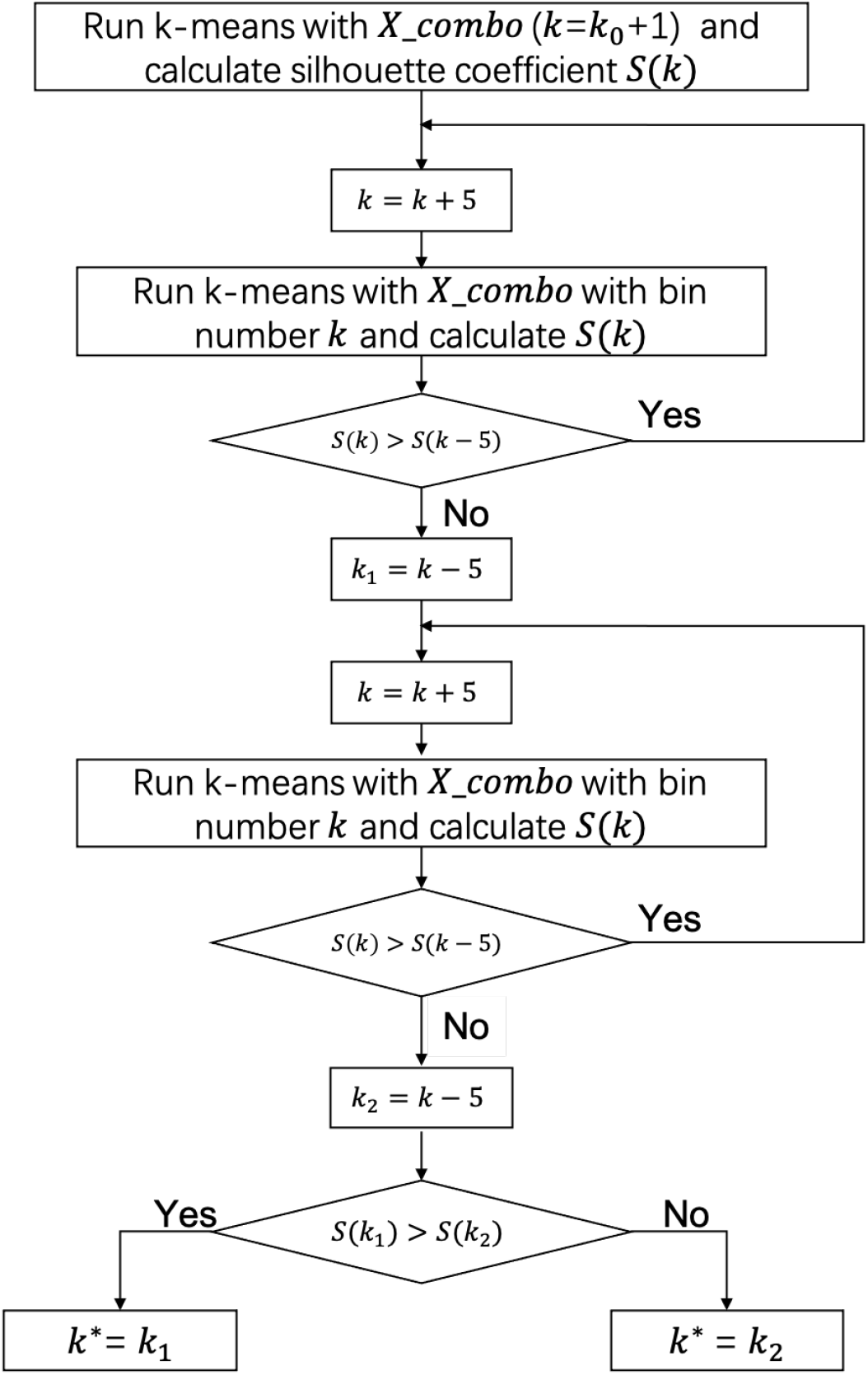
The workflow of estimating the number of bins from *k*_0_.

**Table A1.** The total number of predicted bins per binner and the true bin number for each dataset.

| Methods | CAMI Airways | CAMI Gastrointestinal tract | CAMI Mouse gut | STEC |
| --- | --- | --- | --- | --- |
| MaxBin | 437 | 205 | 867* | 303 |
| MetaBAT | 211 | 159 | 592* | 218 |
| CONCOCT | 202 | 155 | 344* | 331 |
| VAMB | 344 | 217 | 641 | 256 |
| BMC3C | 1560 | 342 | 565 | 921 |
| DAS Tool | 137 | 156 | 577* | 140 |
| MetaWRAP | 144 | 139 | 523 | 155 |
| Metabinner | 285 | 202 | 641 | 194 |
| True bin number (contigs>1,000 bp) | 754 | 246 | 769 | ... |

The results with "\*" are from the CAMI tutorial (Fernando Meyer, et al. 2021, Nature protocols).

**Table A2.** Performance comparison of the binners on the real dataset evaluated by CheckM.

| Dataset | Methods | Metrics |  |  |  |  |  |
| --- | --- | --- | --- | --- | --- | --- | --- |
|  |  | #bins<br>(>50%<br>comp<br><10%<br>cont) | #bins<br>(>70%<br>comp<br><10%<br>cont) | #bins<br>(>90%<br>comp<br><10%<br>cont) | #bins<br>(>50%<br>comp<br><5%<br>cont) | #bins<br>(>70%<br>comp<br><5%<br>cont) | #bins<br>(>90%<br>comp<br><5%<br>cont) |
| STEC (metaSPAdes assembly) | CONCOCT | 101 | 72 | 22 | 92 | 65 | 21 |
|  | MaxBin | 102 | 72 | 28 | 86 | 61 | 26 |
|  | MetaBAT | 91 | 53 | 18 | 87 | 50 | 18 |
|  | VAMB | 129 | 81 | 25 | 127 | 79 | 25 |
|  | BMC3C | 108 | 65 | 21 | 107 | 65 | 21 |
|  | DAS Tool | 78 | 57 | 24 | 73 | 52 | 24 |
|  | MetaWRAP | 143 | 96 | 31 | 135 | 92 | 31 |
|  | MetaBinner | **147** | **101** | **33** | **141** | **96** | **32** |

The best results among all the methods are in bold, while the best results among the individual binners are italicized. The input binning results of MetaWRAP are generated by CONCOCT, MaxBin and MetaBAT. "#bins (>50% comp <10% cont)" denotes that the number of recovered bins that have >50% completeness and <10% contamination.

where *D*(*x*) is the shortest distance between point *x* and the chosen initial centers.

## Availability of data and materials

All the simulated datasets used in this study are publicly available from the CAMI website (https://data.cami-challenge.org). The assembled contigs of the STEC dataset are available from

https://drive.google.com/file/d/1QISGgQNre5Cqut3x9NENiLN1wLmGICyP/view?usp=sharing. Source codes for MetaBinner are freely available at the https://github.com/ziyewang/MetaBinner.

## Ethics approval and consent to participate

Not applicable.

## Consent for publication

All authors have approved the manuscript for submission.

## Authors’ contributions

SZ conceived and supervised the project. SZ and ZW designed the study and the methodological framework. ZW and RY implemented the methods. PH and ZW carried out the computational analyses. ZW drafted the paper. FS and SZ modified the paper. FS, SZ and ZW finalized the paper. All authors agree to the content of the final paper.

